# Protein preference and elevated plasma FGF21 induced by dietary protein restriction is similar in both male and female mice

**DOI:** 10.1101/2022.04.22.489137

**Authors:** K.L. Volcko, J.E. McCutcheon

## Abstract

Animals that are moderately protein restricted respond to this dietary stress by increasing consumption of protein-containing foods. This is true in many species, including rodents. Rodent models of protein restriction have typically relied on only male subjects, and there are plausible reasons why female rodents may respond differently to dietary protein restriction. To address this gap in knowledge, the current experiments examined protein preference after two weeks on a 5% protein diet or 20% protein control diet, in male and female mice. We found that female mice, like male mice, increase consumption of 4% casein (protein) relative to 4% maltodextrin (carbohydrate) when presented with both simultaneously. Interestingly, this increased consumption was due to more bursts in females and more licks per burst in males, indicating possible differences in mechanism by which increased intake is achieved. Stage of the estrous cycle did not affect female responses. Moreover, we measured plasma FGF21 – a hormone induced by protein restriction and necessary for protein preference – in male and female mice. Here, we found no statistical differences between protein-restricted males, females in diestrus, or females in proestrus. In non-restricted mice FGF21 levels were low, but significantly higher in females in proestrus than females in diestrus or males. Overall, these experiments highlight the importance of including female subjects in studies of food choice and macronutrient restriction.

**Highlights:**

- Protein-restricted male and female mice exhibit protein preference
- Estrous cycle stage does not influence protein preference
- Estrous cycle stage does not affect plasma FGF21 during protein restriction
- Non-restricted females in proestrus have higher FGF21 than those in diestrus or males

## 1. Introduction

It is of huge importance to be able to sense current nutritional needs, and to direct behavior towards seeking and consuming certain indispensable nutrients. Of the three macronutrients, the intake of protein is most tightly regulated. Protein cannot be stored in the body for future use, and many amino acids can only be obtained through the diet. Protein restriction affects many aspects of metabolism and behavior. Behavioral adaptations to protein restriction include an increased appetite for foods containing protein. This is true in species ranging from male and female humans to fruit flies (Liu et al., 2017; Ribeiro & Dickson, 2010; Vargas et al., 2010), as well as in male rodents (Chiacchierini et al., 2021; Hill et al., 2019; Murphy et al., 2018). Alarmingly, behavioral responses to protein restriction have been vastly understudied in female rodents, despite the evidence for sex differences in protein requirements (Didelija et al., 2017; Leibowitz et al., 1991) and in metabolic responses to both protein restriction (Larson et al., 2017) and restriction of certain amino acids (Forney et al., 2020; Jonsson et al., 2021). As rodent research is crucial for understanding the neurophysiology of protein appetite, this represents a major gap in knowledge.

Males and females show many differences in eating (reviewed in (Asarian & Geary, 2013), some of which are mediated by ovarian hormones. Estradiol, for example, profoundly affects ingestive behavior, typically by suppressing intake (Czaja et al., 1983; Santollo et al., 2013), but has also recently been discovered to increase intake in times of need (e.g. for food or water) (Santollo et al., 2021; Yu et al., 2020). This novel role for estradiol in controlling food intake is intriguing when viewed through the lens of protein homeostasis. It seems plausible that estradiol could act to increase appetite for protein when dietary protein is scarce, but this hypothesis has not yet been tested. One mechanism by which estradiol could influence protein appetite is through fibroblast growth factor 21 (FGF21), which signals low protein status to the brain (Chaumontet et al., 2019; Hill et al., 2019; Laeger et al., 2014, 2016; Martin et al., 2021) and which is necessary for protein preference in male mice (Hill et al., 2019). FGF21 is increased by protein restriction in males and in ovariectomized females, and to a lesser degree in females when cycling normally (Larson et al., 2017). This difference between intact and ovariectomized females indicates that ovarian hormones likely influence FGF21 levels in a protein-restricted state. Indeed, estradiol has been shown to induce FGF21 transcription (Allard et al., 2019; Badakhshi et al., 2021; Hua et al., 2018) and plasma FGF21 is higher in non-restricted female mice in proestrus than estrus or diestrus (Hua et al., 2018). It is not yet known if these estradiol-associated differences in FGF21 cause corresponding changes in protein appetite.

The purpose of the present study was to determine if protein-restricted female mice show a preference for protein similar to that seen in protein-restricted male mice. Moreover, we sought to examine more closely if sex and/or cycle stage affects FGF21 levels, and if FGF21 levels are correlated with protein preference.

## 2. Material and methods

### 2.1 Animals

Adult male and female C57BL/6NRj mice (6 – 8 weeks old) were purchased from Janvier (France) and housed in a temperature- and humidity-controlled room, maintained on a 12:12 light:dark cycle (lights-on at 7:00). Mice were purchased and tested in two cohorts; cohort 1 was all males (n = 16) and group- or single-housed. Cohort 2 included both males (n = 16) and females (n = 32). In cohort 2, two same-sex mice shared each cage, separated by a perforated divider that allowed visual, olfactory, and auditory communication but prevented physical contact. Water and food were available *ad libitum*. Half the mice were fed a control diet (20% casein; D11051801; Research Diets) and the other half an isocaloric protein-restricted diet (5% casein; D15100602; Research Diets). Mice were maintained on the diets for at least one week before any behavioral testing began. Body weight was measured approximately 5 d per week in cohort 1. Food intake, body weight, and female cycle stage was measured daily in cohort 2. Cycle was monitored by squirting 20 μl of saline into the vagina, pipetting up and down 2-3 times, and expelling the fluid onto a glass slide. Slides were visually examined and the appearance of the fluid was used to decide the day of testing and plasma collection. All animal care and experimentation were in compliance with the EU directive 2010/63/EU for animal experiments.

### 2.2 Testing apparatus

Testing took place in operant chambers (24 cm × 20 cm; Med Associates, Fairfax, VT) equipped with two bottles connected to contact lickometers. Both bottles were located on the same wall, approximately 11 cm apart. Sippers were accessible through a circular hole, 20 mm in diameter. When only one bottle was present, the hole for the missing sipper was plugged. The fan and house light were on continuously during testing. Hardware was controlled and the onset and offset of each lick were recorded using MEDPC-V software (Med Associates).

### 2.3 Behavioral test of protein preference

Protein preference was assessed with a paradigm adapted from that used previously by our laboratory with rats (Murphy et al., 2018). Testing took place in operant chambers during the light phase, between approximately 8:00 – 15:00. Mice were habituated to drinking in the chambers for 1 h per day for three days. On the first two days a single bottle of 0.2 mM sucralose (69293, Sigma) was available (alternating sides), and on the third day bottles of sucralose were available on both sides. Mice were then, over 4 days, given alternating daily access to a carbohydrate solution (4% maltodextrin; 419672, Sigma) and a protein solution (4% casein; C8654, Sigma, with 0.21% methionine; M9625, Sigma) for 24 h in the home-cage. Both solutions included 0.2 mM sucralose and 0.05% Kool-Aid (grape or cherry). Presentation of each solution (carbohydrate on day 1 and 3, protein on day 2 and 4 and vice versa) and flavor (grape paired with protein, cherry paired with carbohydrate and vice versa) were counterbalanced. After the home-cage access, mice were given a 1 h session in the operant chambers on each of two days, with each solution (maltodextrin and casein) presented a single time on different sides of the chamber. Next, protein preference was assessed in a 1 h session in the operant chamber. During this test, both maltodextrin and casein solutions were presented simultaneously, on the same sides as during the single-bottle sessions. Females were tested for protein preference on either proestrus or diestrus. Mice had been on the experimental diets for 16 – 19 days at the time of the preference test (Fig. 1).

**Figure 1.**
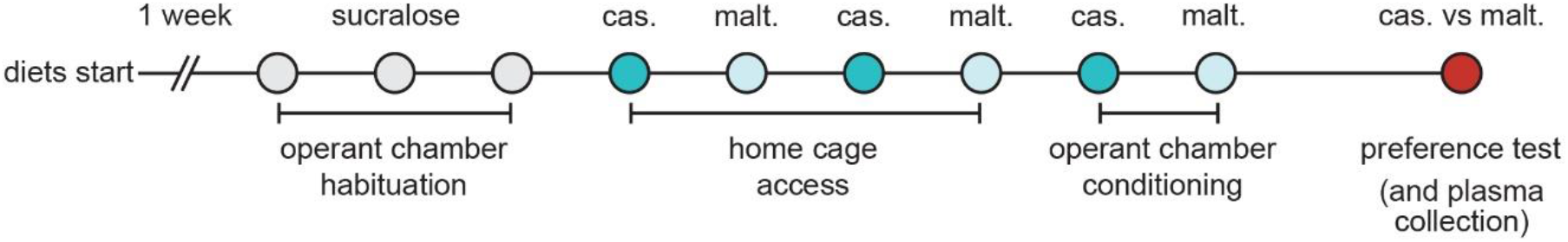
Schematic of the experimental design. Mice were given at least one week on the experimental diets before any behavioral testing began. After three days of habituation to drinking sucralose in operant chambers, mice were given home cage access to casein (cas.) or maltodextrin (malt.) on alternating days. Mice were then given two sessions in the operant chamber, one with a bottle of casein available and one with a bottle of maltodextrin available. Finally, mice were tested for protein preference in a two-bottle choice test in the operant chamber and blood collected 3 – 6 h later.

### 2.4 Plasma levels of FGF21

Female mice and male mice in cohort 2 were killed by decapitation under isoflurane anesthesia on the day of the protein preference test (approximately between 17:00 – 18:00). Trunk blood was collected in K2 EDTA-coated tubes (16.444.100, Sarstedt) and placed on ice. Blood was then centrifuged, within an hour of collection, for 10 minutes at 3000 rpm and 4°C (Mikro 220R centrifuge). Plasma was separated and frozen at −70°C until analysis. FGF21 was assessed by ELISA (Millipore #EZRMFGF21-26K) on a ClarioStar microplate reader.

### 2.5 Data analysis and availability

Licks were recorded and used as a proxy for intake. Recording individual licks also allowed us to assess drinking microstructure by examining interlick intervals. A burst was defined as series of at least 3 licks with <1 s interlick intervals.

Due to the hypotheses regarding the role of sex and cycle stage on protein preferences, we conducted separate statistical analyses on male mice and on female mice. All statistics were conducted using SPSS (IBM Corp, Armonk, NY, USA, version 28.0.1.0).

Body weight was analyzed with repeated-measures ANOVA with Time as within-subjects factor and Diet as between-subjects factor. Percent change in body weight was analyzed as a one-way ANOVA with Diet as factor. Average food intake for 8 of the 9 days between diet initiation and start of home-cage nutrient access was analyzed by one-way ANOVA with Diet as factor. Change in food intake and change in body weight from previous day, were analyzed by repeated measures ANOVA with Cycle Stage as within-subjects factor and Diet as between-subjects factor. Total Licks in the One-Bottle Conditioning Sessions was analyzed by repeated-measures ANOVA with Nutrient as within-subjects factor and Diet as between-subjects factor. Total Licks, Burst Size, and Burst Number in the Two-Bottle Preference Test were analyzed similarly, with the exception that Cycle Stage was an additional within-subjects factor for female mice. FGF21 levels were analyzed by one-way ANOVA with Cycle Stage (male, female-diestrus, and female-proestrus) as factors; NR and PR were split for these analyses. Undetectably low ELISA results were coded as 0 for subsequent analyses. Significant interactions were further probed with pair-wise comparisons and Bonferroni-corrected to account for multiple comparisons.

All data and custom analysis scripts are available at the following DOI: https://doi.org/10.5281/zenodo.6420813. Some individual data points are missing due to technical issues.

## 3. Results

### 3.1 Food intake, body weight, and cycle stage

Body weight increased from start of diet to the day of the preference test in both female (F_1, 30_ = 84.991, p < 0.001) and male (F_1, 30_ = 18.951, p < 0.001) mice (Fig. 2A). We did not, however, find a main effect of Diet in either females (F_1, 30_ = 0.005, p = 0.947) or males (F_1,30_ = 0.585, p = 0.451), nor a Diet by Time interaction in females (F_1,30_ = 2.552, p = 0.121) or males (F_1, 30_ =0.923, p = 0.344). Similarly, percent body weight change from diet start to preference test did not differ between control and protein-restricted female (F_1,31_ = 2.540, p = 0.121) or male (F_1,31_ = 0.924, p = 0.344) mice (Fig. 2B).

**Figure 2.**
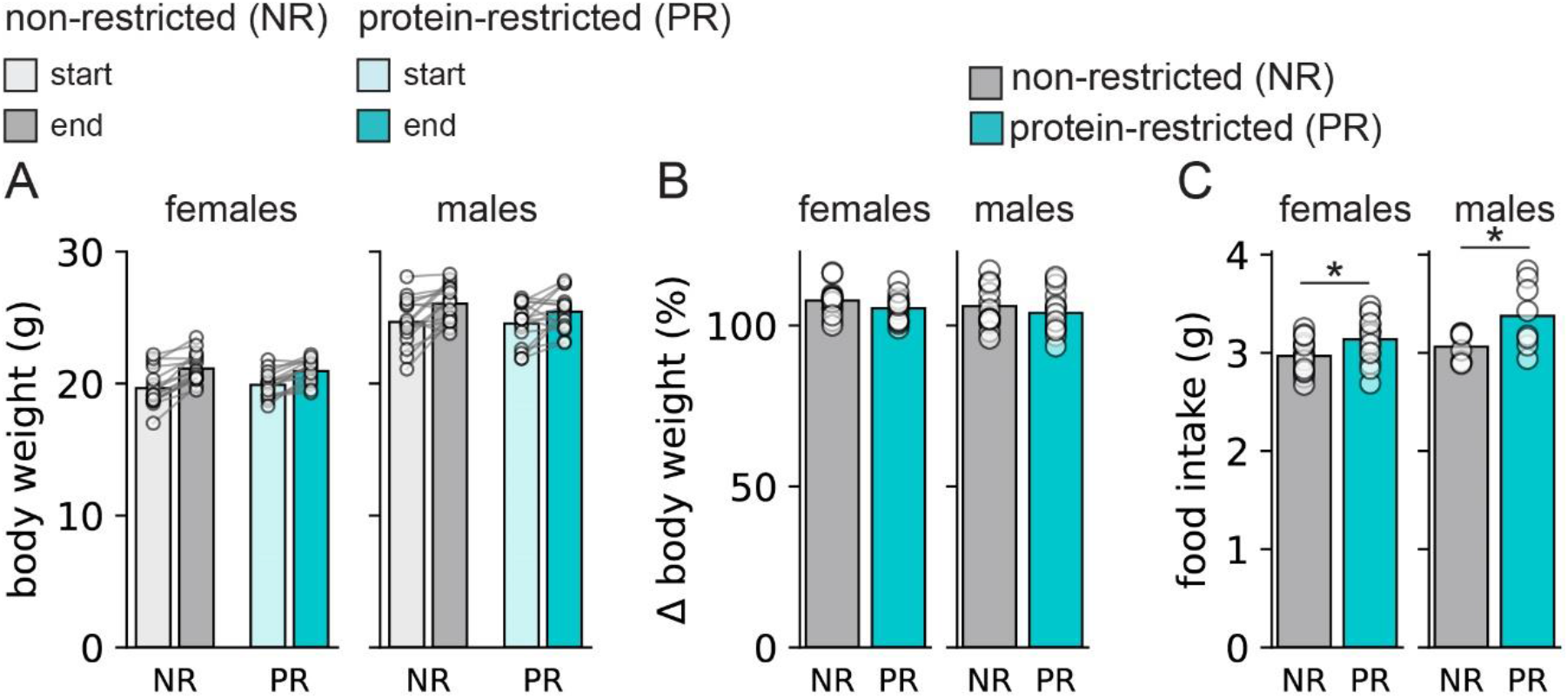
Changes in body weight and food intake. Non-restricted and protein-restricted female and male mice gained weight between the start of the experimental diet and the day of the preference test (A), and percent change in weight gain was the same for all groups (B). Food intake was greater in protein-restricted mice of both sexes than in non-restricted controls (C). Bars are mean and circles show data from individual mice. * p < 0.05 vs. non-restricted mice of the same sex.

Food intake was measured in females and a subset of males (cohort 2). Average daily intake for 8 of the first 9 d on the experimental diets, from day 1 until the home-cage nutrient access, was significantly higher in protein-restricted female (F_1, 31_ = 6.039, p = 0.020) and male (F_1, 15_ = 5.787, p = 0.031) mice than in non-restricted controls (Fig. 2C).

Visual examination of vaginal discharge revealed stable 4–5-day cycles in the majority of female mice. Female mice were tested for protein preference, and subsequently killed for blood collection, on diestrus or proestrus. We compared daily change in food intake and body weight across two cycles for each mouse. We found no main effect of Diet for either change in food intake (F_1,30_ = 0.030, p = 0.864) or change in body weight (F_1,30_ = 0.692, p = 0.412), nor a Diet by Cycle Stage interaction for change in food intake (F_1,30_ = 0.061, p = 0.807) or change in body weight (F_1, 30_ = 0.004, p = 0.949). There was a significant main effect of Cycle Stage for both change in body weight (F_1,30_ = 38.249, p < 0.001) and change in food intake (F_1,30_ = 19.218, p < 0.001). Pairwise comparisons between estrus and proestrus, estrus and diestrus, and proestrus and diestrus were therefore conducted for both change in body weight and change in food intake. Body weight change did not differ between diestrus and proestrus (p = 0.486) but was significantly lower on estrus than on either diestrus (p < 0.001) or proestrus (p < 0.001) (Fig. 3A). Similarly, change in food intake did not differ between diestrus and proestrus (p = 0.249) but was significantly lower on estrus than on either diestrus (p < 0.001) or proestrus (p = 0.0116) (Fig. 3B).

**Figure 3.**
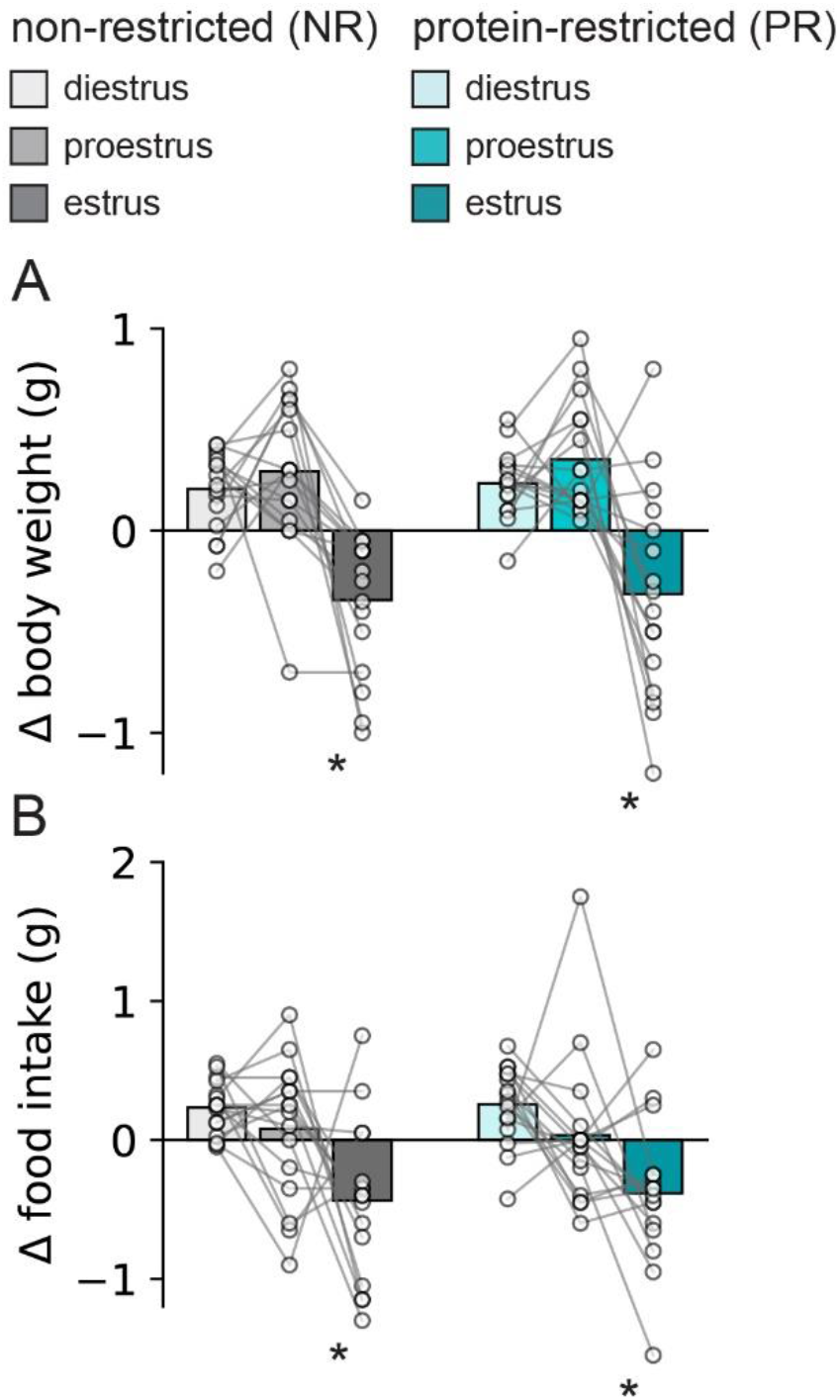
Change in body weight and food intake across the estrous cycle. Estrous cycle stage was measured by vaginal swabs and confirmed post hoc by examining change in food intake and body weight. Females in both diet groups showed the expected reduction in body weight (A) and food intake (B) on estrus compared to on diestrus or proestrus. Bars are mean and circles show data from individual mice. * p < 0.05 vs. non-restricted mice of the same sex.

### 3.2 One-Bottle Conditioning Sessions

Licks for maltodextrin and licks for casein were measured during 1-bottle operant chamber conditioning sessions when either casein or maltodextrin solution was available. We found no main effects of Nutrient (F_1,28_ = 0.629, p = 0.434) or Diet (F_1,28_ = 0.546 p = 0.466) in males, though there was a significant Nutrient by Diet interaction (F_1,28_ = 4.914, p = 0.035). Pair-wise comparisons, however, did not show significant differences between maltodextrin and casein in each diet group. Similarly, in females we found no main effects of Nutrient (F_1,28_ = 1.852, p = 0.184) or Diet (F_1,28_ = 0.088 p = 0.769), and only a weak trend towards an interaction (F_1, 28_ = 3.684, p = 0.065). On the whole, the results of the 1-bottle conditioning sessions indicate that intake of maltodextrin and intake of casein were approximately the same in non-restricted and protein-restricted mice (Fig. 4).

**Figure 4.**
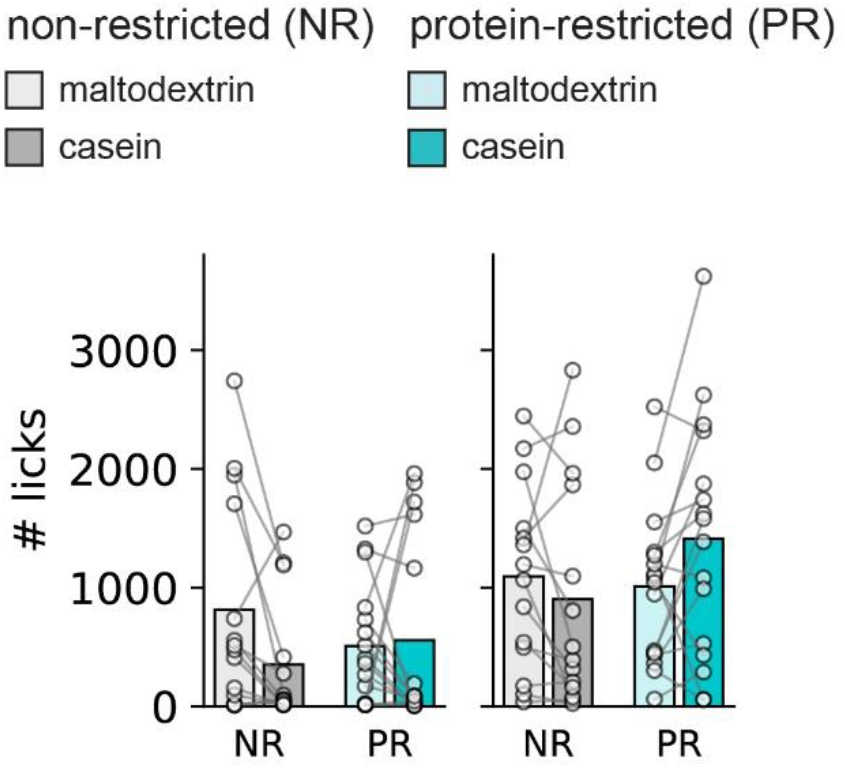
No difference in nutrient intake in one-bottle conditioning sessions. Non-restricted (NR) and protein-restricted (PR) mice received a single bottle of either maltodextrin or casein on each of two days. Intake was equivalent for both nutrients in each diet group of both sexes. Bars are mean and circles show data from individual mice.

### 3.3 Two-Bottle Preference Test

Following 1-bottle conditioning sessions, mice had access to both casein and maltodextrin solutions in the same session while licking behavior was monitored (Fig. 5A). Cumulative intake suggested higher casein intake in PR than NR mice (Fig. 5B), and total intake over the course of the session was compared between diet groups of each sex. In females there were no main effects of Nutrient (F_1,28_ = 4.049, p = 0.054), Diet (F_1,28_ = 2.005, p = 0.168), or Cycle Stage (F_1,28_ = 0.003, p = 0.959). There was, however, a significant Nutrient x Diet interaction (Fig. 5C; F_1,28_ = 12.993, p = 0.001), and pairwise comparisons between licks for maltodextrin and licks for casein within each diet group showed no difference in non-restricted females (p = 0.521) but significantly more licks for casein than maltodextrin in protein-restricted females (p = 0.004). Protein preference (ratio of casein-to-total intake) was also significantly higher in protein-restricted than non-restricted females (Fig. 5D; F_1,31_ = 10.164, p = 0.004), and Cycle Stage did not influence protein preference (F_1,31_ = 1.111, p = 0.301). We also tested for a difference from 50% preference in each diet group. Non-restricted females did not have a protein preference different from 50% (t_15_ = 1.547, p = 0.143) while protein-restricted females showed a greater than 50% preference for casein (t_15_ = 3.233, p = 0.006). To gain greater insight into potential reasons for differences in intake, we examined the microstructure of drinking. Burst size (i.e. licks per burst) showed a significant Nutrient by Diet interaction (F_1,28_ = 6.852, p = 0.014), but pairwise comparisons between licks for maltodextrin and licks for casein within each diet group showed no difference in NR (p = 0.31) or PR (p = 0.08) females (Fig. 5E). There was a trend towards a main effect of Cycle Stage for burst size (F_1, 28_ = 2.06, p = 0.050), with mice in proestrus trending towards fewer licks per burst. Burst number also had a significant Nutrient by Diet interaction (Fig. 5F; F_1, 28_ = 11.542, p = 0.002), with pairwise comparisons revealing no difference in burst number to casein or maltodextrin in NR females (p = 0.16) but significantly more bursts to casein than maltodextrin in PR females (p = 0.006). Although no cycle stage differences were found, data also displayed broken down by cycle stage for preference test number of licks (Fig. 6A), burst size (Fig. 6B), and burst number (Fig. 6C).

**Figure 5.**
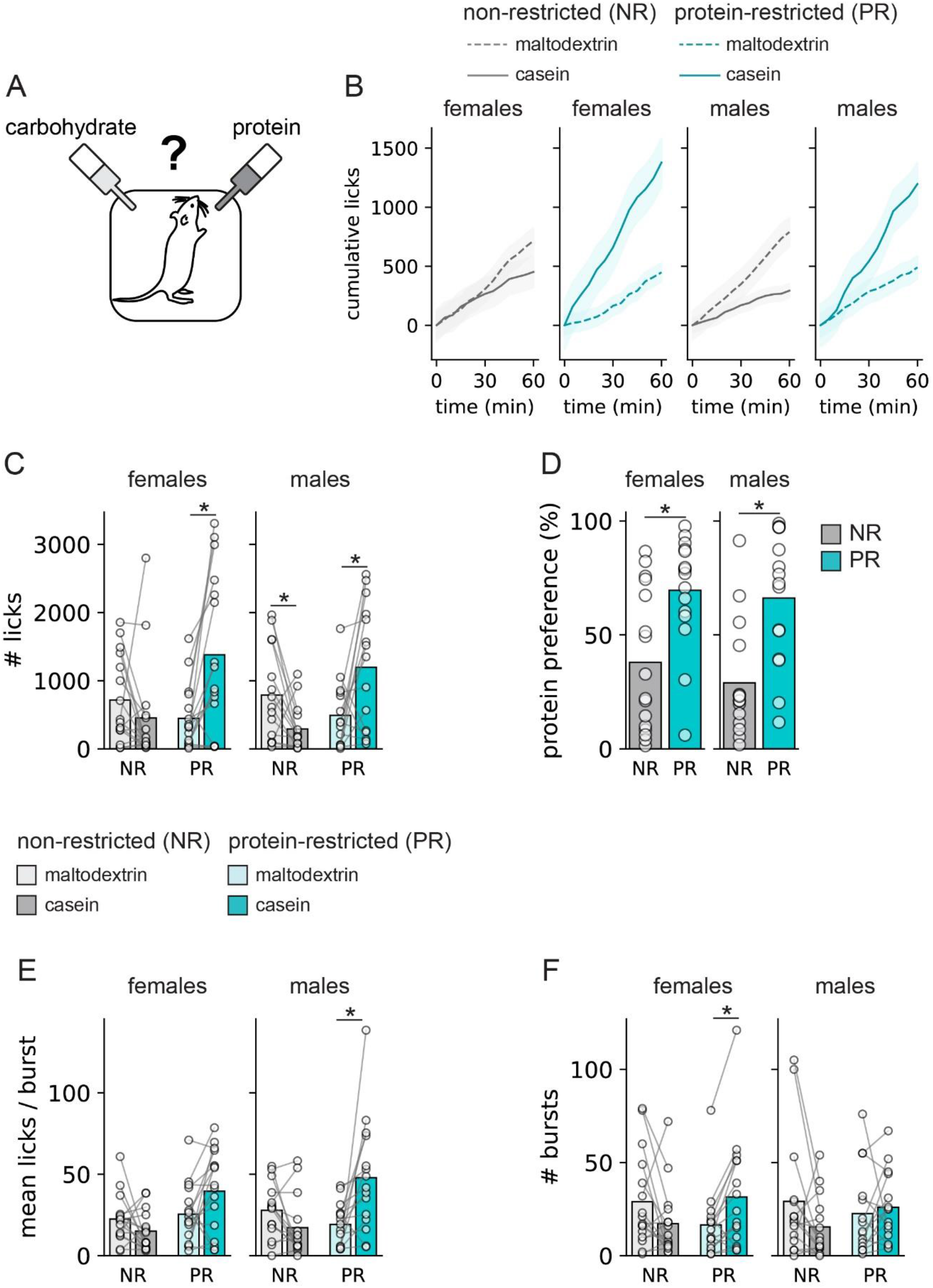
Protein preference in male and female mice in two-bottle test. Mice were allowed to freely choose between two bottles, one containing a carbohydrate solution and the other a protein solution (A). Cumulative licks for the nutrients (maltodextrin, dashed line; casein, solid line) in female and male non-restricted control and protein-restricted mice (B). Protein-restricted (PR) male and female mice both licked more to protein than carbohydrate, while non-restricted (NR) male mice licked more to carbohydrate than protein (C). Protein preference was elevated in both PR male and female mice (D). Analysis of lick microstructure showed that burst size (mean licks per burst; E) and number of bursts (F) differed by sex with burst size for casein, relative to maltodextrin, increasing in PR males and number of bursts for casein, relative to maltodextrin, increasing in PR females. Bars are mean and circles show data from individual mice. * p < 0.05 vs. maltodextrin in the same diet group (C, E, F) or vs. non-restricted mice of the same sex (D).

**Figure 6.**
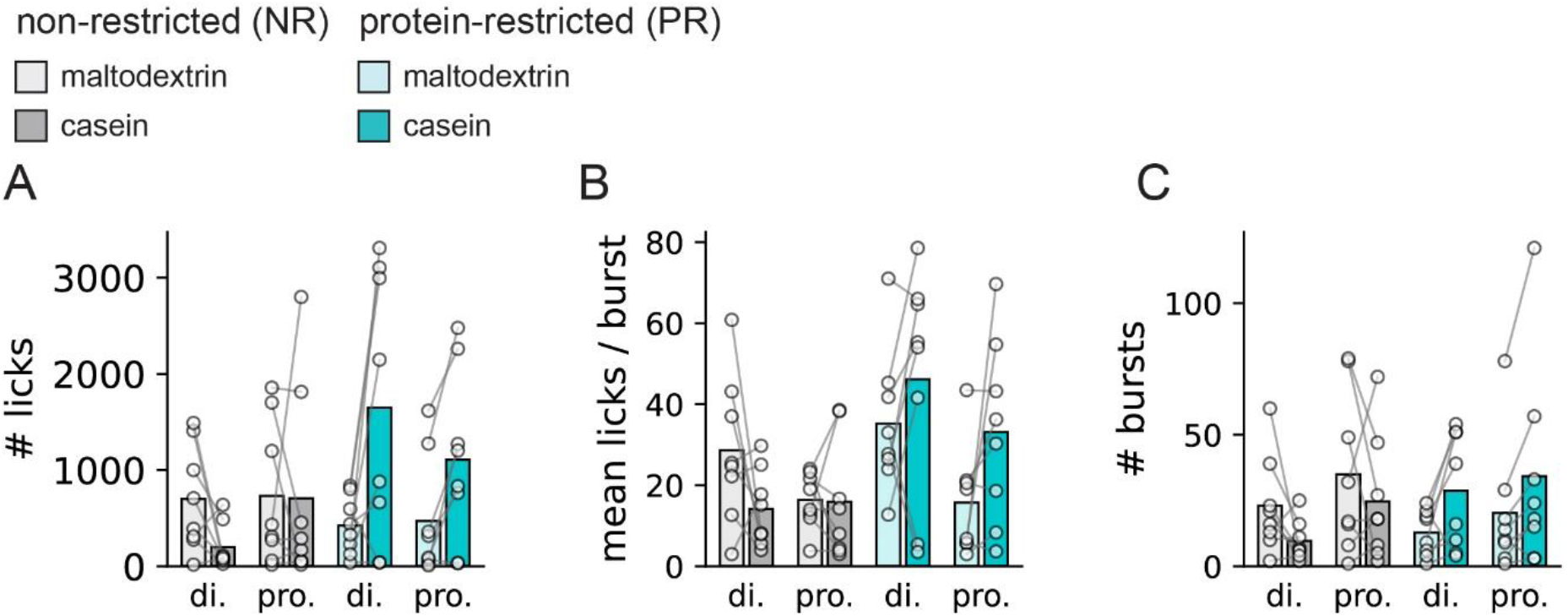
Cycle stage does not affect behavior in the two-bottle preference test. No significant differences were found in total licks (A), mean licks per burst (B), or burst number (C) when diestrus was compared to proestrus in female mice. Bars are mean and circles show data from individual mice.

In males we found no main effects of Nutrient (F_1, 28_ = 0.888, p = 0.354) or Diet (F_1,28_ = 3.174, p = 0.086), but a significant Nutrient by Diet interaction (Fig. 5C; F_1,28_ = 16.864, p < 0.001). Pairwise comparisons between maltodextrin and casein for each diet group revealed that non-restricted males drank more maltodextrin than they did casein (p = 0.041), while the opposite was true in protein-restricted males, which drank more casein than maltodextrin (p = 0.014). Accordingly, protein preference was higher in protein-restricted than non-restricted male mice (Fig. 5D; F_1, 29_ = 20.824, p < 0.001). Both non-restricted and protein-restricted male rats had a protein preference statistically different from 50%, with non-restricted males having a less than 50% preference (t_14_ = 3.244, p = 0.006) and protein-restricted males having a greater than 50% preference (t_14_ = 3.210, p = 0.006). We also examined drinking microstructure in males during the preference test. Just as in females, males had a significant Nutrient by Diet interaction in burst size (Fig. 5E; F_1,28_ = 12.100, p = 0.002), and in males pairwise comparisons showed no difference in burst size to casein or maltodextrin in NR mice (p = 0.11) but significantly more licks per burst in PR mice (p = 0.026). Burst number, however, had no significant Nutrient by Diet interaction (Fig. 5F; F_1,28_ = 2.851, p = 0.102), nor main effects of either Nutrient (F_1,28_ = 1.053, p = 0.314) or Diet (F_1,28_ = 0.081, p = 0.778).

### 3.4 Plasma FGF21

We collected blood after the preference test, and measured plasma FGF21. We found that several samples (11 out of 24, all from non-restricted mice), had undetectably low levels (below 10 pg/ml). We found a significant effect of Diet on plasma FGF21 (F_1,46_ = 40.875, p < 0.001), such that protein-restricted mice had higher FGF21 than non-restricted controls did, but no effect of Sex (F_1,46_ = 0.279, p = 0.389) nor a Diet by Sex interaction (F_1,46_ = 0.522 p = 0.24). However, we had reason to believe a high degree of variability in FGF21 in female mice due to sampling at two distinct phases in the cycle, and previous reports finding differences between protein-restricted freely-cycling and ovariectomized females (Larson et al., 2017) and non-restricted females at different cycle stages (Hua et al., 2018). Our interest was in understanding how cycle stage may influence FGF21 levels in a protein-restricted state. Therefore, we conducted a one-way ANOVA on FGF21 between PR males, PR females in diestrus, and PR females in proestrus. There was no significant difference between these groups (Fig. 7C; F_2, 23_ = 2.912, p = 0.077). We did, on the other hand, find a significant negative correlation between FGF21 and protein preference in PR mice, such that mice with higher FGF21 had lower protein preference (Fig. 7D; R^2^ = − 0.583, p = 0.004). When PR mice were separated by Cycle Stage, however, this negative correlation was only apparent in females in proestrus (R^2^ = − 0.880 p = 0.004) and absent in females in diestrus (R^2^ = −0.608, p = 0.110) and males (R^2^ = − 0.246 p = 0.557).

**Figure 7.**
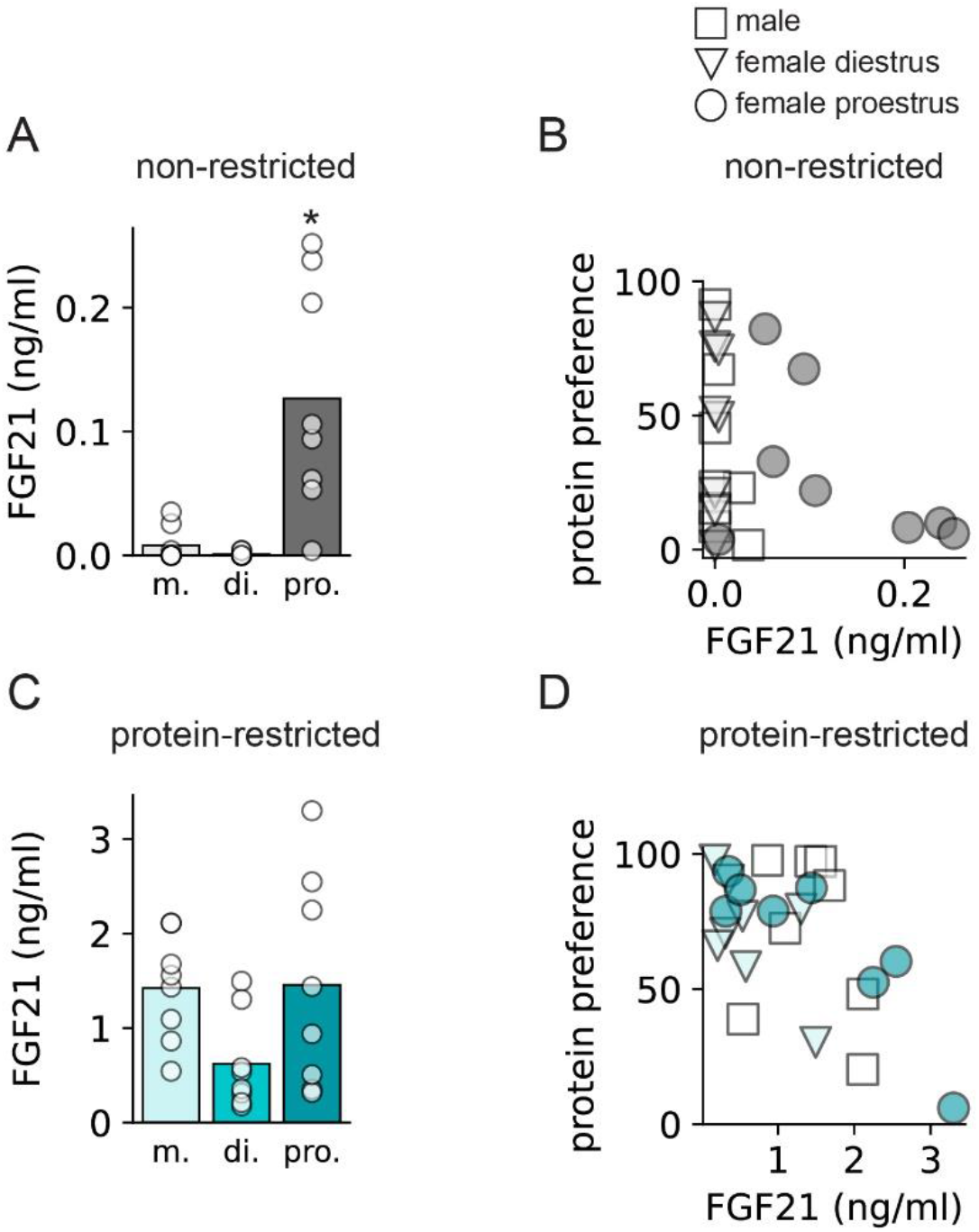
Plasma FGF21 is elevated by protein restriction. Non-restricted (NR) female mice in proestrus had higher levels of FGF21 than did females in diestrus or males (A). There was no correlation between protein preference and FGF21 in NR mice (B) There were no differences in plasma FGF21 in protein-restricted males, females in diestrus, or females in proestrus (C), but there was a negative correlation between protein preference and FGF21 in PR mice in proestrus (D). In A and C, bars are mean and circles show data from individual mice. In B and D, each square, triangle, or circle is an individual mouse * p < 0.05 vs. all other groups.

Although our primary interest was in FGF21 during protein restriction, we also compared FGF21 in NR mice in a separate analysis. In this case, we found that NR males, NR females in diestrus, and NR females in proestrus had significantly different plasma FGF21 levels (Fig. 7A; F_2, 22_ = 12.542 p < 0.001). Bonferroni-corrected pair-wise comparisons indicated that females in proestrus had higher FGF21 than those in diestrus (p = 0.006), or males (p = 0.009). Females in diestrus and males did not differ from one another (p = 0.53). In NR mice there was no correlation between FGF21 and protein preference (Fig. 7B; R^2^ = −0.325, p = 0.121) when analyzed as a single group The high number of non-detectable samples in NR males and NR females in diestrus made correlations between FGF21 and protein preference hard to run in these groups, but NR females in proestrus did not have a significant correlation between the two variables (R^2^ = −0.477, p = p = 0.523).

## 4. Discussion

The present experiments have given us several important insights into the role of sex in protein appetite. First, and perhaps most important, is the finding that protein-restricted female mice develop a preference for a protein solution over a carbohydrate solution in much the same manner as males do (Hill et al., 2019; Murphy et al., 2018).The fact that female mice exhibit protein preference is not surprising in and of itself, as females of other species have been shown to shift their preference towards protein when protein-restricted (Liu et al., 2017; Ribeiro & Dickson, 2010; Vargas et al., 2010). Rather, it is of note that the same paradigm leads to such similar behavior in both male and female mice. There are known sex differences in other features of protein consumption. For example, when allowed to choose freely between three macronutrient diets, protein makes up a greater percentage of daily intake in male than in female rats (Leibowitz et al., 1991), and restriction of the amino acid arginine is better tolerated by female than male mice (Didelija et al., 2017). This suggest that males may require more protein than females do, and that the same low-protein diet for the same amount of time could represent more severe protein restriction for males than for females. Thus, we anticipated that females might require a different length of time on the protein-restricted diet, a different concentration of protein in the diet, or different conditioning parameters (e.g. concentration of nutrients, number of sessions), in order to show protein preference. However, we did not find evidence for this. Whether such differences exist under other conditions, such as sooner after the diet manipulation begins, remains to be seen. We did find a sex difference in microstructure of drinking, in which female mice drank more casein than maltodextrin due to changes in burst number, while male mice drank more because of changes in burst size. These subtle changes imply that increased protein consumption during times of need may be achieved by quite different means in male and female mice. Changes in burst size indicate changes in orosensory feedback, while changes in burst number indicate changes in postingestive feedback (reviewed in (Smith, 2001)). Our findings therefore suggest that male mice found the protein solution more palatable than the carbohydrate solution, while female mice drank more of the protein solution because it was perceived as less satiating than the carbohydrate solution. In rats, studies of microstructure when drinking casein and maltodextrin solutions found both increased burst number and burst size in the protein-restricted state (Murphy et al., 2018). The differences in findings between the present experiment and that by Murphy et al. (2018) could be due to species differences, or different criteria for burst thresholds.

Second, we found that protein preference was not affected by cycle stage. The role of cycle stage in protein consumption in a non-restricted state is not clear. Some studies have found that rats show no difference in percent of diet made up by protein across the cycle (Abadie et al., 1993), while others have found that protein makes up a larger proportion of intake at estrus (Wurtman & Baum, 1980). Whether or not cycle stage affects protein consumption in a non-restricted state does not necessarily mean that the same would be true in a restricted state. Although the estrous cycle involves fluctuations of many systems, one of the major hormones with a cyclic pattern of release is estradiol. Dietary amino acids regulate liver estrogen receptor alpha activity (Della Torre et al., 2011), showing that estradiol may be sensitive to protein intake. Moreover, estradiol has different effects on intake, depending on the need state of the animal (Santollo et al., 2021; Yu et al., 2020). As such, one hypothesis is that estradiol may promote protein appetite during protein restriction, leading to higher protein preference at proestrus than at diestrus. Here, however, we did not find differences in protein preference at the different cycle stages. Two caveats prevent us from drawing conclusions about the specific role of estradiol in protein appetite: 1) we did not measure estradiol directly, and 2) protein preference was high in both groups, creating a possibility of ceiling effects. Future experiments should address these limitations.

Third, we found that plasma FGF21 and protein preference were inversely correlated in protein-restricted mice. This is somewhat surprising, given the necessity of FGF21 for protein preference (Hill et al., 2019). This implies one of two things: 1) that although some amount of FGF21 is needed to induce protein preference, the relationship between the two is not straightforward and may be an inverted-U rather than linear, or 2) the increased consumption of protein that leads to a high protein preference score lowers plasma FGF21. The fact that the correlation was only found in females in proestrus is interesting, and might imply that females in this cycle stage are especially responsive to protein ingestion. Contrary to our expectations, we did not find a difference in plasma FGF21 between protein-restricted male mice, female mice in proestrus, and female mice in diestrus. Others have found that protein-restricted male mice have nearly twice as high circulating FGF21 as protein-restricted, freely-cycling female mice do, and that ovariectomized females have intermediate levels (Larson et al., 2017). One explanation could be that the freely cycling mice were sampled at a point other than proestrus (e.g., three quarters of their cycle). Furthermore, FGF21 is released in response to many different stressors (for example, (Luo & McKeehan, 2013)) and ovariectomy has far-ranging effects beyond simply eliminating estradiol.

Although we did not find statistically significant differences in FGF21 across cycle stage in protein-restricted mice (p = 0.077), there was high variability, which may have obscured an underlying pattern. This high variability could have several causes, for example, blood was sampled after a series of nutrient conditioning sessions and a preference test, in which individual mice drank different amounts of each nutrient solution. Indeed, as mentioned above, it is possible that the negative correlation between protein preference and FGF21 was due to increased consumption affecting plasma levels of FGF21. To firmly establish the role of cycle in protein-restriction-induced FGF21 levels, plasma should be collected at proestrus and diestrus in protein-restricted females without access to protein solutions. Future experiments should also examine FGF receptor expression in relevant brain regions, across estrous cycle and correlated with protein preference score.

In non-restricted mice, females in proestrus had significantly higher plasma FGF21 than either males or females in diestrus. This is consistent with previous findings that FGF21 is elevated in proestrus relative to the rest of the cycle (Hua et al., 2018), and the stimulatory role that estradiol has on FGF21 (Allard et al., 2019; Badakhshi et al., 2021; Hua et al., 2018). It is interesting that plasma FGF21 is higher in females in proestrus than it is in males. More relevant to our interests, however, was the finding that this difference in FGF21 did not appear to influence protein preference. Protein preference did not differ by cycle stage, and in non-restricted mice there was no correlation between FGF21 and protein preference. This, together with the data from the protein-restricted mice, strongly suggests that there is a threshold above which FGF21 generates a preference for protein over carbohydrate; below this threshold, higher or lower levels do not affect protein appetite, and above the threshold, further elevations do not increase preference.

Estrous cycle can be disrupted by dietary stress. Restriction of protein in particular appears important, as 40% calorie restriction – with all macronutrients equally restricted – causes greater rates of anestrous than 40% calorie restriction when the caloric deficit comes only from fat and carbohydrate (Della Torre et al., 2011). Although some have found that low-protein diets disrupt the estrous cycle in rats (Guilbert & Goss, 1932), we found no evidence that two weeks of a 5% casein diet affected the estrous cycle in mice. Cycle length was regular and consistent in both diet groups. This level of protein restriction, while mild enough to spare reproductive function, nonetheless triggers compensatory behaviors to avoid further restriction when a protein source is available.

In conclusion, we have found that female mice display a similar behavioral response to dietary protein restriction as male mice do, although the reason for increased protein intake (e.g. post-ingestive or orosensory feedback) may differ by sex. Moreover, we found no differences between protein-restricted male and female mice in plasma FGF21 levels. In non-restricted mice, females in proestrus have significantly higher levels of FGF21 than females in diestrus or males – but they do not display higher protein preference. This indicates that below a certain threshold, FGF21 does not induce protein appetite. On the whole, the results from the present experiments serve as an important first demonstration of protein preference in protein-restricted female mice and add to our knowledge about the phenomenon of protein appetite.

## Acknowledgements

The authors wish to thank the excellent animal care staff at UiT; Neoma Boardman for use of a centrifuge and advice on plasma preparation; the UiT Advanced Microscopy Core Facility, and especially Kenneth Bowitz Larsen, for technical assistance; and Destiny Brakey for advice on data interpretation.

